# ZenReg: A modular Python platform for fast and memory-efficient N-dimensional microscopy image registration

**DOI:** 10.64898/2026.08.07.743572

**Authors:** Fabrizio Musacchio, Martin Fuhrmann

**Affiliations:** German Center for Neurodegenerative Diseases (DZNE), 53127 Bonn, Germany

**Keywords:** microscopy image registration, bioimage analysis, multiphoton microscopy, motion correction, phase correlation, NoRMCorre, rigid registration, memory mapping, reproducibility, FAIR data

## Abstract

Motion artifacts are almost unavoidable in functional time-lapse and structural volumetric multiphoton microscopy. They arise from respiration, heartbeat, locomotion, awake behavior, instrument heating, mechanical vibration, and slow drift, while the recorded signal is often photon-limited, blurred by scattering, and biologically time varying. Consequently, motion correction is frequently an essential prerequisite for quantitative bioimage analysis rather than a merely cosmetic preprocessing operation. Edge- and landmark-centric registration strategies are often poorly matched to these data because useful structures may be sparse, diffuse, out-of-focus, or changing in fluorescence intensity. We present ZenReg, an open-source Python platform that formulates common 2D+t, 3D, and 3D+t microscopy registration tasks as modular, geometry-preserving alignment problems. ZenReg combines FFT- and intensity-based translational registration, projection-based rotation estimation, piecewise translational motion correction, and dense or sparse six-degree-of-freedom volume registration within one canonical microscopy stack model. Disk-backed arrays support chunked processing of large or remote image stacks, and every run can produce registered images together with shift tables, correlation metrics, summary plots, and machine-readable settings.

In synthetic benchmarks with known ground truth, ZenReg recovered global 2D and 3D translations with subpixel accuracy across moderate noise and drift regimes. High-noise and large-drift tests separated the registration models: FFT-based methods failed abruptly once image information or shared support became insufficient, intensity-based translational alignment degraded more gradually under severe noise, and piecewise translational correction improved spatially varying local motion where a single global transform was inadequate. In real biological data, ZenReg increased mean template correlation in a 3000-frame calcium-imaging movie from 0.334 to 0.487 and recovered imposed continuous three-photon volume motion with a mean translational error of 0.055 px or, for six-degree-of-freedom rigid motion, a mean shift error of 0.068 px and a mean rotation error of 0.015 degrees.

By coupling modular registration backends to transparent sidecar outputs, ZenReg turns motion correction into an inspectable, memory-aware, and FAIR-oriented component of reproducible bioimage analysis.

## Introduction

Quantitative bioimage analysis depends on spatial consistency. In time-lapse microscopy, the same biological structure is commonly sampled repeatedly, but the observed intensity pattern is displaced by instrument drift, sample motion, scanning distortions, or animal movement. If left uncorrected, these displacements contaminate downstream measurements such as fluorescence traces, structural colocalization, object morphology, volumetric occupancy, and longitudinal comparisons. Motion correction is therefore not a cosmetic preprocessing step: it determines whether a measured temporal change can be interpreted biologically or is merely the consequence of moving the sample relative to the acquisition coordinate system.

This problem is particularly important in multiphoton microscopy. Two-photon microscopy made cellular and subcellular functional imaging possible in intact tissue, but awake or behaving subjects introduce motion artifacts that can be fast, nonuniform, and correlated with the behavior under study [1]. Because raster-scanning microscopes sample pixels sequentially rather than instantaneously, within-frame motion can distort the apparent shape of the field of view. Axial motion is harder still: lateral displacement can often be corrected by aligning frames, whereas movement out of the focal plane changes the recorded fluorescence intensity and can no longer be treated as a simple in-plane translation [1]. Recent work on axial and real-time three-dimensional motion correction further illustrates that volumetric and functional multiphoton imaging remain tightly constrained by movement, photon budget, and acquisition speed [2]. Three-photon microscopy improves deep-tissue access and enables chronic or deep in vivo imaging, including through-skull and prefrontal cortex applications, but it does not remove the computational challenge: deep volumes can be dim, scattering remains substantial, and repeated volumetric acquisitions create large arrays whose registration must be both accurate and practical [3, 4].

The image statistics of these data differ from those of many anatomical radiology volumes. In magnetic resonance imaging or computed tomography, tissue interfaces, organs, and modality-specific anatomical boundaries often provide information for feature-, surface-, intensity-, or mutual-information-based alignment [5, 6]. In multiphoton microscopy, the relevant signal can instead consist of sparse puncta, blurred cell bodies, diffuse neuropil, weak structural markers, or activity-dependent fluorescence changes [7, 8, 9]. Edges are often ambiguous, contrast varies over time, and the same cell may brighten or dim for biological reasons. For this reason, the safest registration models in routine microscopy are often geometry-preserving: translations, rotations, or full rigid transformations. Affine scaling and shearing can be mathematically convenient, but for structural microscopy they may alter biological morphology and therefore become scientifically undesirable unless explicitly justified.

Correlation-based translational registration is consequently a natural baseline. Phase correlation, based on the Fourier shift theorem and the normalized cross-power spectrum, estimates image displacement from the location of a correlation peak in Fourier space [10, 11]. In practice, these Fourier-domain operations are evaluated efficiently using the fast Fourier transform (FFT), and we therefore refer to this family as FFT-based registration methods. Phase correlation is fast, robust to many intensity-scale changes, and extends from two-dimensional images to three-dimensional volumes. Other registration families complement this baseline when the motion model or image statistics require it: multiresolution intensity-based schemes estimate subpixel transforms by minimizing image differences [12], NoRMCorre estimates piecewise translational motion fields from overlapping spatial patches to approximate nonuniform motion in calcium imaging [13], and full-volume rigid optimizers estimate rotations and translations in physical 3D space [14, 15].

These methods are supported by a mature open-source ecosystem, but the tools emphasize different layers of the microscopy registration problem. ImageJ/Fiji and ImageJ2 provide broad plugin-based environments for biological image analysis [16, 17]. The StackReg/TurboReg family provides established intensity-based subpixel registration models [12]. scikit-image provides reusable Python image-processing algorithms, including phase-correlation registration primitives [18]. CaImAn and Suite2p place motion correction inside larger calcium-imaging pipelines that also perform source extraction or ROI analysis [19, 9]. SimpleITK, ITK, elastix, and ANTs-style workflows provide powerful image-registration engines, especially for medical and anatomical images [14, 15, 20]. Thus, many algorithmic components already exist, but they are not usually presented as one registration-specific workflow for heterogeneous microscopy files, channel-coupled correction, metadata propagation, batch processing, and shareable quality-control outputs. Table 1 summarizes this landscape by design emphasis. The comparison does not attempt to enumerate every extension that an expert user could assemble manually. Instead, it highlights which capabilities are available as primary end-to-end workflow features. The gap is practical as much as algorithmic. Some tools provide strong registration models but are embedded in larger analysis suites. Others expose flexible primitives but leave microscopy-specific axis handling, metadata propagation, reporting, and batch reproducibility to the user. General medical-image registration frameworks provide powerful 3D optimization engines, but their abstractions are not centered on time-lapse microscopy stacks, channel propagation, or common 2D+t and 3D+t workflows. Meanwhile, the data itself has grown large enough that the computational representation matters: a 20–40 GB stack cannot be casually duplicated in memory several times during registration, and files stored on a server should not be repeatedly reread during parameter tuning.

**Table 1:** Representative position of ZenReg within the open-source registration and bioimage-analysis ecosystem. “Partial” indicates that a capability can often be assembled through plugins, scripting, or external workflows, but is not the primary end-to-end emphasis of the cited tool.

| Tool | Primary emphasis | 2D+t | 3D+t | Full 3D rigid | Memory-aware | Report side-cars | Microscopy workflow role |
| --- | --- | --- | --- | --- | --- | --- | --- |
| ImageJ/Fiji / StackReg | Interactive bioimage analysis and plugin registration | Yes | Partial | Partial | Partial | Partial | Broad environment; workflows depend on plugins and macros |
| scikit-image | Algorithm library | Yes | Yes | Partial | No | No | Reusable primitives, not a complete microscopy workflow |
| CaImAn / NoRMCorre | Calcium-imaging analysis suite | Yes | Partial | No | Yes | Partial | Strong calcium-imaging pipeline with source extraction |
| Suite2p | Calcium-imaging pipeline | Yes | Partial | No | Yes | Partial | Fast functional-imaging pipeline centered on ROI extraction |
| SimpleITK / ITK | Medical and general image registration engine | Partial | Yes | Yes | Partial | No | Powerful optimizer backend, not microscopy-stack workflow layer |
| elastix | Intensity-based registration framework | Partial | Yes | Yes | Partial | No | Parameterized registration engine for broad image-registration problems |
| ZenReg | Microscopy registration platform | Yes | Yes | Yes | Yes | Yes | Unified registration wrapper, TZ-CYX axes, metadata propagation, and reproducible outputs |

ZenReg was developed to address this combined mathematical and infrastructural problem. It is not meant to replace every specialized registration package. Instead, it provides a single, inspectable platform for common microscopy registration models, all expressed on the same canonical image representation, with memory-efficient execution, parallelizable work units, and reproducible outputs. The scientific contribution is therefore twofold. First, ZenReg unifies several geometry-preserving registration models that are directly relevant to 2D+t, 3D, and 3D+t microscopy. Second, it treats registration as a FAIR-oriented analysis step: the registered image is written together with the estimated motion parameters, similarity metrics, plots, and settings needed to understand and rerun the computation [21].

## Methods

ZenReg was designed by treating registration as a linked mathematical and practical problem. The mathematical problem is to estimate a restricted, geometry-preserving transformation between a moving microscopy observation and a reference. The practical problem is to make that estimate usable in everyday laboratory workflows, where files are multidimensional, metadata-rich, often large, and commonly explored by changing parameters over several attempts. The following sections therefore first define the registration model, then describe the design goals and architecture that connect this model to executable workflows, and finally introduce the implemented registration families and result-assessment outputs.

### Registration problem

Let a microscopy experiment be represented as a non-negative image array

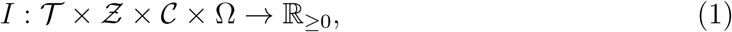

where *T* indexes time, *Z* indexes axial planes, *C* indexes acquisition channels, and Ω *⊆* Z^2^ denotes the lateral pixel grid. A single time point and channel is therefore a volume

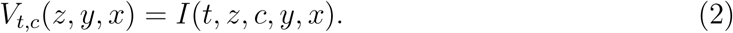

For two-dimensional time series, *Z* has cardinality one. The registration task is to estimate, for each moving time point *t*, a transformation Φ*_t_* that aligns the selected registration signal *V_t,c∗_* to a reference *R*, while applying the same estimated transformation to all channels:

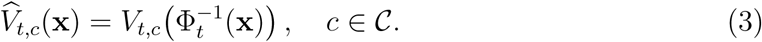

This channel-coupled formulation is important in microscopy because a high-contrast structural or anatomical channel can be used for motion estimation while functional or marker channels are carried along without estimating independent, physically impossible channel-specific motion.

ZenReg restricts the default transformation family to geometry-preserving models. The simplest model is a translation

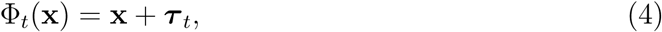

where ***τ****_t_* = (*τ_y_, τ_x_*) for two-dimensional registration and ***τ*** *_t_* = (*τ_z_, τ_y_, τ_x_*) for full-volume registration. For rigid-volume registration, the model becomes

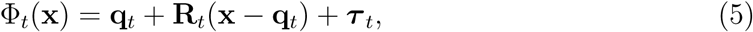

where **R***_t_ ∈ SO*(3) is parameterized by rotations around the *z*, *y*, and *x* axes, and **q***_t_* is the rotation center. No scaling or shearing is included by default, because those degrees of freedom can change biological morphology and are therefore inappropriate for many structural microscopy questions.

The reference *R* may be a selected frame, a projected volume, or a template derived from multiple aligned frames. In all cases, the estimate is obtained by maximizing a similarity functional

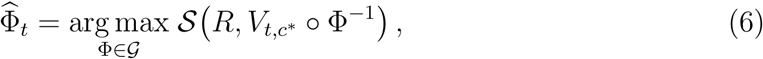

where *G* is the chosen transformation family and *S* is a correlation, phase-correlation, mean-squared-difference, or information-theoretic metric depending on the backend. For anisotropic volumes, the physical spacing vector **s** = (*s_z_, s_y_, s_x_*) is part of the geometry, because a rotation in voxel coordinates is not equivalent to a rotation in physical space unless *s_z_* = *s_y_* = *s_x_*.

### Design goals and architecture

The implementation follows five design goals, summarized in Box 1. These goals are not independent software preferences, they are the operational constraints that follow from the registration problem in microscopy. A correction should preserve geometry unless a stronger model is scientifically justified, should be estimated in a predictable coordinate system, should remain practical for large files, and should leave enough evidence behind to be reviewed and reproduced. ZenReg’s architecture is a direct response to these requirements.

The high-level workflow in Figure 1 starts with the canonical stack model. Input and output are delegated to OMIO, which normalizes supported microscopy inputs to time–z–channel–y–x axes and returns metadata with shape, axis, physical-size, channel, and timing information [22, 23, 24, 25]. This makes ZenReg format-agnostic at the registration level: users can work with supported microscopy file types without manually pre-formatting them into a package-specific array layout. Once the data are represented as *T × Z × C × Y × X*, the same reference selection, channel propagation, projection, and reporting logic can be applied to 2D+t, 3D, and 3D+t data.

**Figure 1:**
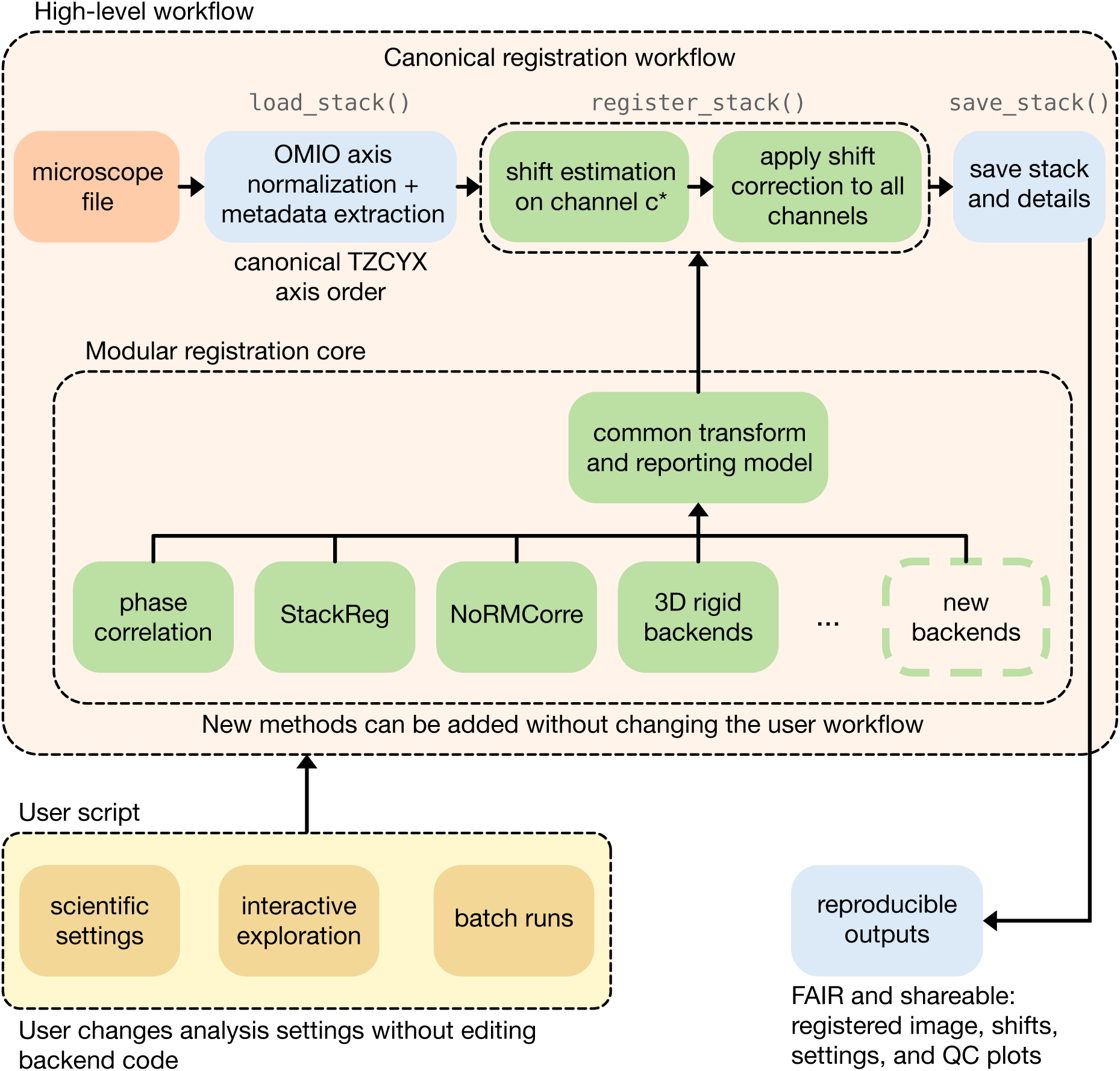
ZenReg architecture and registration workflow. ZenReg is built as a modular registration platform: file I/O, axis normalization, motion estimation, transform application, reporting, and user interaction are separated into exchangeable layers with shared data and transform conventions. Heterogeneous microscope files enter through “load_stack”, where OMIO normalizes axes and extracts metadata into the canonical time–z–channel–y–x (“TZCYX”) representation. “register_stack” estimates motion from a selected registration channel and applies the resulting correction to all channels. Phase correlation, StackReg, NoRMCorre, full 3D rigid backends, and future methods connect to a common transform and reporting model, so new algorithms can be added without changing the user workflow. “save_stack” then writes registered images together with reproducible, shareable outputs such as shift tables, settings, metadata, and quality-control plots. User scripts remain deliberately separate from the core implementation: they expose scientific settings for interactive exploration or batch processing without requiring users to edit backend code.

#### Box 1: Design goals of *ZenReg*

1. **Geometry-preserving registration for microscopy.** Routine workflows should emphasize translations and rigid transformations before more deformable models, so that biological structures are not changed by the correction itself.
2. **One canonical representation across file formats.** Registration should operate on a predictable time, axial, channel, and lateral axis order while preserving microscopy metadata whenever possible.
3. **Memory-efficient processing of large and remote stacks.** A stack should not need to fit completely in RAM after loading, and repeated parameter tuning should not require repeated full reads from slow network storage.
4. **A small user-facing workflow surface.** Users should be able to perform standard analyses by specifying scientific settings rather than rewriting registration logic.
5. **Reproducible, shareable outputs.** Motion parameters, similarity metrics, settings, and quality-control plots should be written next to the registered image so that a registration result can be audited and reused.

Within the modular registration core, as illustrated in the inner box of Figure 1, each backend estimates a transformation in its own way, but all backends return motion parameters in the same coordinate convention and all transformations are applied channel-wise through a common resampling path. This addresses the first design goal by keeping translations and rigid transformations explicit, and it addresses extensibility by allowing new estimators to be added without changing the user-facing workflow. Phase correlation, StackReg-style intensity fitting, NoRMCorre-style patch fields, projection-based rotation, dense 3D rigid optimization, and sparse point-based rigid alignment are therefore backends behind a shared microscopy registration model rather than isolated scripts.

As indicated in Figure 1, ZenReg deliberately separates the high-level workflow and user-interaction layer from the tested core implementation. A typical user script specifies the input path, reference time point, registration channel, transformation family, shift limits, interpolation, and memory settings, while the core performs validation, registration, channel propagation, optional cropping, metadata updates, and reporting. This separation keeps analysis scripts readable enough for interactive work and documentation, but prevents users from needing to edit the lower-level implementation in order to run ordinary analyses. It also makes registered outputs easier to compare across projects, because the same core writes the same sidecar formats for different workflows.

Memory-aware execution is a separate layer used when needed rather than a different registration method (Figure 8a). Disk-backed Zarr caches can be requested through OMIO for large or remote files. ZenReg then processes the slices, projections, time points, or volumes required by the selected model and can write registered output to a disk-backed result array. Parallel work is distributed over independent time points or axial slices where the mathematical model permits it. Full 3D rigid registration necessarily touches whole volumes, but projection-based translation, slice-wise intra-stack correction, and many output operations remain chunk-compatible.

### Projection-based templates

For volumetric stacks, many fast microscopy workflows estimate lateral motion from projections rather than from full volumes. ZenReg formalizes this as an operator

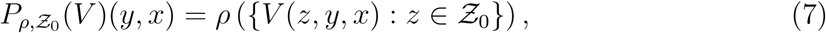

where *Z*_0_ is an optional axial subrange and *ρ* is a projection statistic. Maximum projections emphasize sparse puncta and bright landmarks; means preserve dense extended signal; medians reduce outlier influence; variance and standard deviation projections emphasize contrast-rich structure rather than absolute intensity. Projection-based registration is efficient because a 3D+t task can be reduced to a sequence of two-dimensional registrations, after which the same estimated lateral correction is applied to every slice in the corresponding volume.

This approximation is appropriate when motion is dominated by lateral displacement and the axial content is stable enough for a projection to represent the same anatomy over time. It is less appropriate when axial shifts are large, when z-dependent structures enter or leave the projection range, or when full three-dimensional rotations are present. In those cases, ZenReg can estimate full-volume translations or full rigid-volume transformations.

### Fourier phase-correlation translation

For translational registration, ZenReg uses phase correlation as the default estimator. Let *f* and *g* be a fixed and moving image, and suppose that *g*(**x**) *≈ f* (**x** *−* ***δ***). Their Fourier transforms satisfy the shift relation

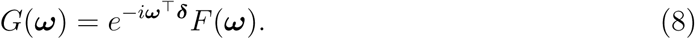

The normalized cross-power spectrum is

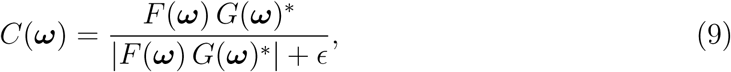

and the inverse transform of *C* contains a peak at the displacement. Subpixel refinement follows efficient local upsampling around the peak [11]. In ZenReg this estimator is used for two-dimensional time series, for projected 3D+t volumes, and for full 3D translations when volumetric registration is selected.

When full 3D translation is unnecessary, ZenReg can still estimate an axial component by applying the same phase-correlation principle to orthogonal projections. Lateral motion is estimated from the axial projection, while axial motion is inferred from projections onto planes that contain the z axis. This projection-based axial estimate is faster than full-volume registration but less complete than directly estimating a displacement in 3D Fourier space.

### Intensity-based StackReg-style registration

ZenReg also supports StackReg-style intensity registration, based on multiresolution minimization of image-intensity differences [12]. In this family of methods, a moving image is transformed by a parameterized model Φ***_θ_*** and fitted by minimizing

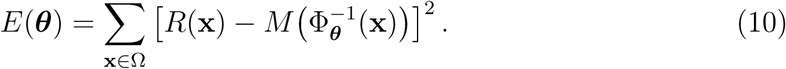

The multiresolution pyramid improves convergence by first fitting coarse image representations and then refining the transform at higher resolution. In ZenReg this route is used as a familiar alternative for two-dimensional and projection-based microscopy workflows.

### NoRMCorre-style piecewise translation fields

To approximate nonuniform in-plane motion without introducing arbitrary affine deformations of the whole image, ZenReg implements a NoRMCorre-style piecewise-rigid model [13]. The field of view is divided into overlapping patches Ω*_k_*. For each patch, a local translation ***δ****_t,k_* is estimated against a template, and the patch shifts are interpolated into a smooth dense displacement field

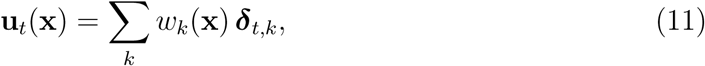

where *w_k_* are spatial interpolation weights. The corrected image is then

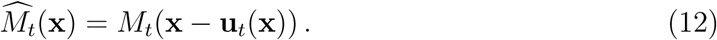

Template construction is an essential part of this model. ZenReg follows the batch NoRMCorre idea of estimating motion against a template that can be updated from corrected frames, including median-of-chunk updates that reduce the influence of transient fluorescence events. In 3D mode the same idea is applied to local volumes rather than planar patches.

The model should be interpreted as a smooth field of translations, not as a general rotation or deformation engine. Small rotations can sometimes be approximated locally, especially near the center of rotation, but large rotations, perspective changes, and complex tissue warping require explicit rigid or nonrigid models. This distinction is important for interpreting NoRMCorre outputs in benchmarks.

### Rotation and full rigid-volume registration

For two-dimensional rotation or rotation around the optical axis of a volume, ZenReg estimates an in-plane angle by transforming fixed and moving projections into polar coordinates. Rotation in Cartesian coordinates becomes translation along the angular axis of the polar image, allowing phase correlation to estimate the angle. Because a rotation can induce residual translation after resampling, ZenReg uses an alternating refinement:

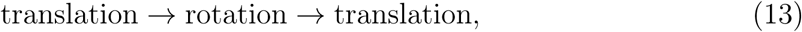

with additional iterations when needed.

For true 3D rotation, ZenReg treats the problem as full rigid-volume registration. The dense backend uses SimpleITK to refine a six-degree-of-freedom Euler transform with multiresolution optimization and either correlation or mutual-information-like metrics [14, 15]. The initial estimate is obtained from orthogonal projections: rotation around *z* from the *xy* projection, rotation around *x* from a *zy* projection, and rotation around *y* from a *zx* projection. The final optimization is performed in physical space, using the voxel spacing derived from microscopy metadata unless manually overridden.

Let **x** denote physical coordinates in the fixed reference volume and let

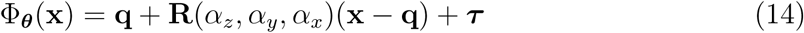

be an Euler rigid transform with center **q**, translation ***τ***, and rotations *α_z_, α_y_, α_x_*. The dense backend solves

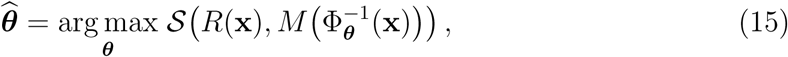

where *S* is normalized correlation for same-modality structural data or an information-theoretic metric for more difficult contrast relationships. The optimization is evaluated in a multiresolution pyramid. If **s** = (*s_z_, s_y_, s_x_*) is the physical voxel spacing, voxel indices **i** are first mapped to physical coordinates **x** = **s** *⊙* **i**. This prevents anisotropic stacks from treating one axial plane as geometrically equivalent to one lateral pixel.

For sparse puncta or spot-like volumes, ZenReg additionally provides a point-based rigid estimator. Peaks are detected in the registration channel, candidate correspondences are found by nearest-neighbor matching, and a rigid transform is estimated from matched point clouds. The optimal least-squares rotation for a matched point set follows the Kabsch solution [26], while robust matching is stabilized by random-sample-consensus-style rejection of inconsistent matches [27]. This backend is intended for puncta-rich structural data where intensity-based volume optimization may be slower or less stable.

Mathematically, let 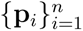 be fixed points and 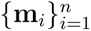 their matched moving points. After subtracting centroids **p̄** and **m̄**, the covariance matrix

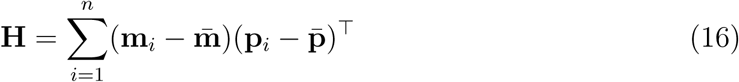

is decomposed by singular value decomposition, **H** = **UΣV***^T^*. The least-squares rigid rotation is

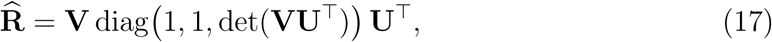

and the translation is ***τ̂*** = **p̄** *−* **R̂m̄**. Random minimal subsets propose candidate transforms. The model with the largest inlier set under a maximum match distance is refined on all inliers. This formulation makes the point backend appropriate when discrete puncta are reliable but global image intensity is a weak registration signal.

### Preprocessing, interpolation, and cropping

Microscopy registration can benefit from light preprocessing for motion estimation. Median filtering suppresses impulse or salt-and-pepper noise before projection or before registration of projected frames [28, 29], while high-pass filtering can suppress slowly varying background in calcium-imaging-like data [13, 19]. ZenReg applies such filters only for estimation unless the user explicitly chooses otherwise. The registered output preserves the original intensity scale.

The resampling step is performed with a selectable interpolation order. Linear interpolation is appropriate for most intensity images; nearest-neighbor interpolation preserves sparse puncta and label-like data; higher-order interpolation can be used when smoothness is preferred. After correction, zero-valued borders may appear because the moving image has been shifted or rotated out of the field of view. ZenReg can crop these borders from the largest detected shifts or from transformed validity masks. For rotations, validity-mask-based cropping is preferable because invalid regions are angled rather than parallel to image borders.

### Assessment and exported results

ZenReg evaluates registration quality by writing estimated translations, rotations, and similarity measures across time. For a fixed reference *R* and corrected moving image 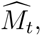, Pearson correlation is computed as

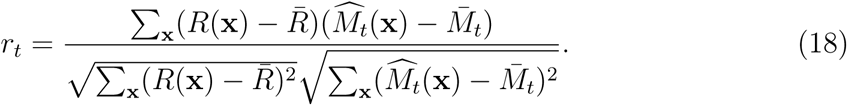

When synthetic ground truth is known, detection accuracy is quantified as absolute error in the correction parameters:

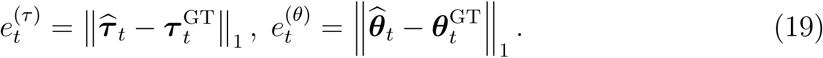

For registered images, residual motion is summarized by changes in frame-to-template correlation and by difference images between reference and moving frames before and after correction. ZenReg writes these quantities as tabular and visual sidecars next to the registered image, together with a settings file that stores the registration model, reference, projection strategy, limits, interpolation, memory mode, and backend settings (Table 2).

**Table 2:**
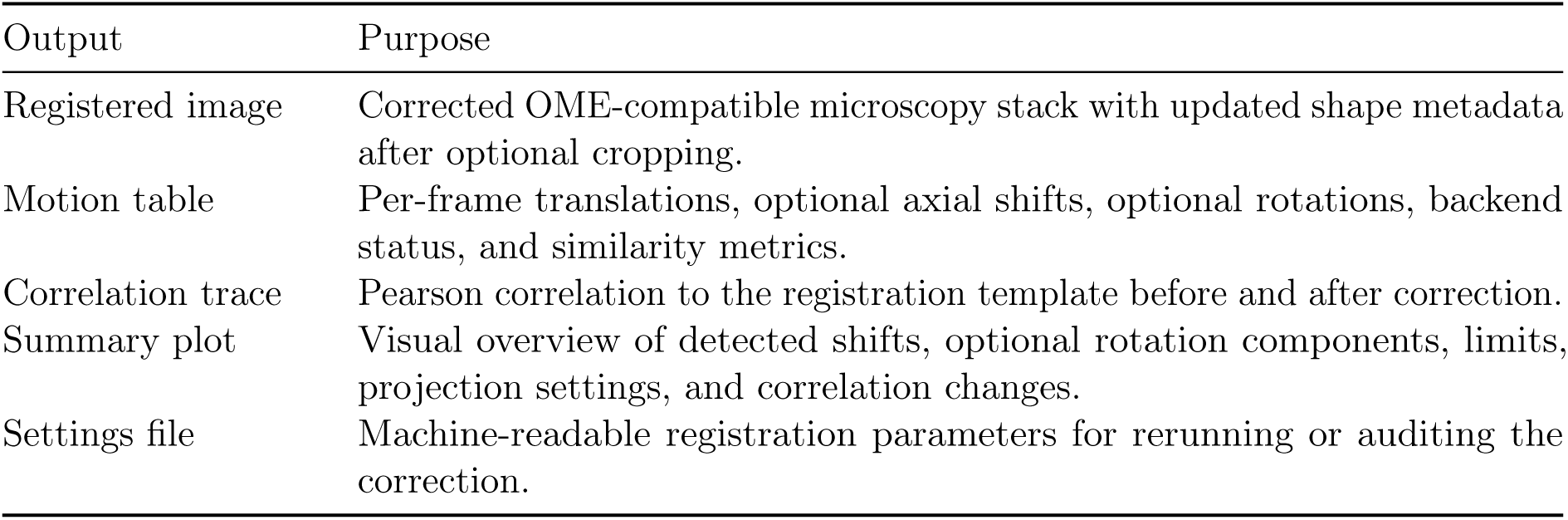
Registration outputs written by ZenReg to make corrected images inspectable and reproducible.

| Output | Purpose |
| --- | --- |
| Registered image | Corrected OME-compatible microscopy stack with updated shape metadata after optional cropping. |
| Motion table | Per-frame translations, optional axial shifts, optional rotations, backend status, and similarity metrics. |
| Correlation trace | Pearson correlation to the registration template before and after correction. |
| Summary plot | Visual overview of detected shifts, optional rotation components, limits, projection settings, and correlation changes. |
| Settings file | Machine-readable registration parameters for rerunning or auditing the correction. |

## Results

### 2D+t registration example

We first evaluated a synthetic two-channel 2D+t benchmark in which 64 time points were translated with known ground-truth motion. Each slice was 256*×*256 pixels. The registration channel contained sparse and moderately blurred structures with additive zero-mean Gaussian noise, while the second channel was transformed by the same ground-truth motion and carried along during correction. Registration used a template aggregated from all time points. For visualization and ground-truth comparison, the estimated correction trace was re-referenced to *t* = 0. This example was chosen to show the complete output logic of ZenReg.

Figure 2a shows the all-frame registration template used for motion estimation. Panels b–d show the raw reference frame (*t* = 0), a representative moving frame (*t* = 25), and their difference image. The red–blue residual pattern in panel d indicates structured displacement: bright blue spots mark reference-frame feature positions where signal is reduced in the moving frame, while bright red spots mark new moving-frame feature positions relative to the reference. After registration, the corresponding panels e–g show that this structured residual is strongly reduced and approaches a noise-dominated difference image. The estimated *y* and *x* corrections follow the ground-truth correction traces across all frames (panel h), template correlation increases after correction (panel i), and per-frame absolute shift error remains near the subpixel range (panel j). Across the 64-frame series, mean template correlation increased from 0.913 before registration to 0.994 after registration, while the mean absolute shift error was 0.126 px and the worst-frame mean error was 0.264 px. Thus, the image-level and parameter-level outputs agree: ZenReg recovers the imposed translational motion and documents the correction quantitatively.

**Figure 2:**
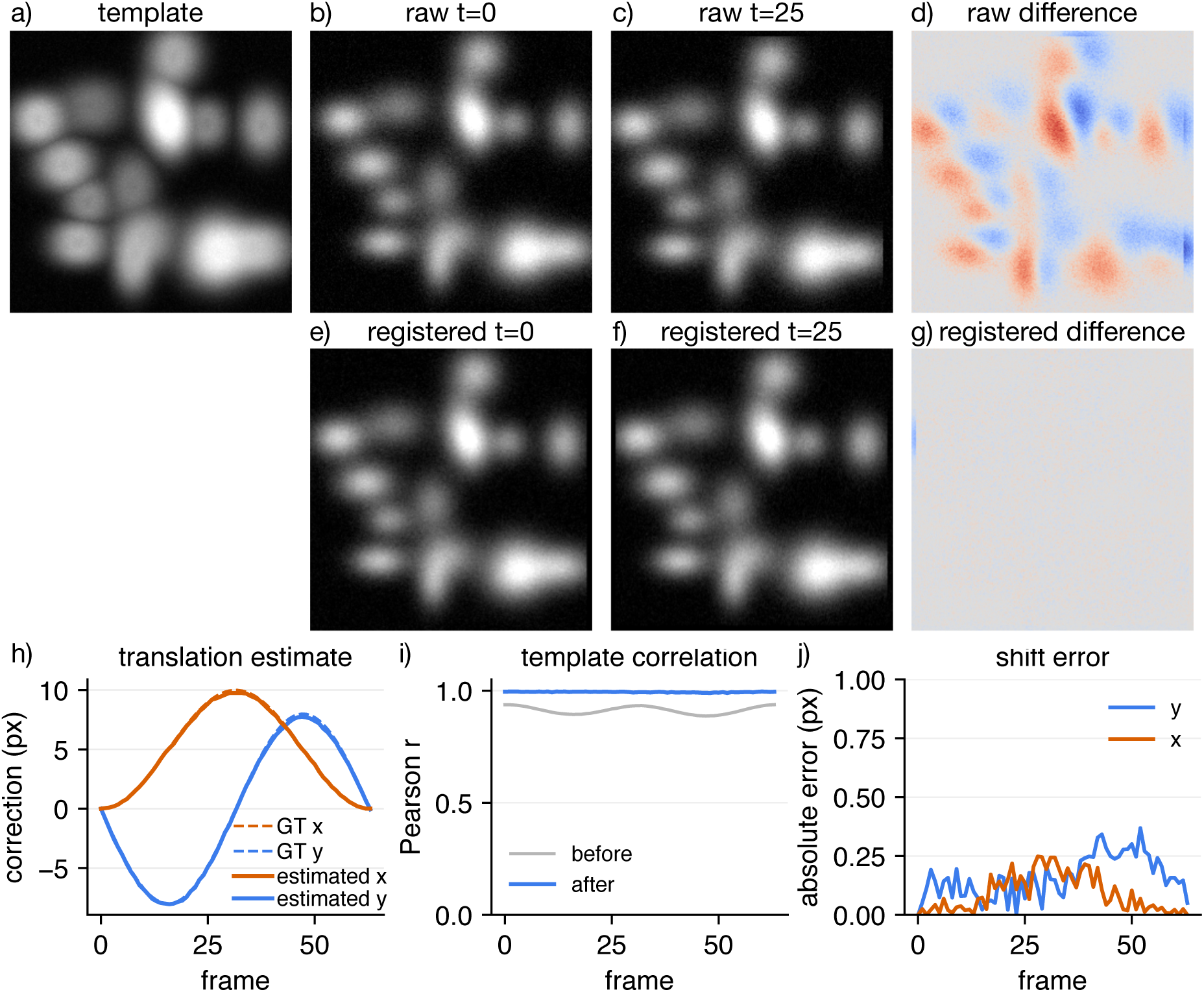
Representative 2D+t registration with known ground truth. The synthetic stack contains 64 time points, one axial plane of 256*×*256 pixels, two channels, moderately blurred spot-like structures in the registration channel, and additive zero-mean Gaussian noise. Registration used an all-frame maximum-intensity template. (a) Registration template. (b–d) Reference frame for visualization (here: *t* = 0), a representative moving frame (*t* = 25), and difference image before correction. (e–g) The same visualization after correction. (h) Detected lateral correction compared with ground truth across time after re-referencing the correction trace to *t* = 0. (i) Pearson correlation to the template before and after correction. (j) Per-frame absolute shift error. The panels are generated by the preprint benchmark and plotting scripts and mirror the quality-control artifacts written by ZenReg for ordinary registrations.

The result also illustrates why ZenReg treats the registered image as only one part of the output. The image panels reveal whether the correction is visually plausible. The motion table reports the estimated parameters. The correlation trace quantifies template similarity before and after correction. And the settings file records how the result was obtained. The registered movie is therefore accompanied by the information needed to audit it.

### Noise and drift benchmarks

We next evaluated how registration accuracy depends on signal-to-noise ratio and drift magnitude. Synthetic 2D+t and 3D+t stacks were generated with known translational motion and increasingly severe noise or displacement amplitudes. We compared phase correlation, StackReg-style intensity alignment, and rigid NoRMCorre-style registration. Separate filtered benchmarks were added for the same regimes: median-filtered projections for phase correlation and StackReg-style alignment, and CaImAn-style spatial high-pass filtering for NoRMCorre.

Figure 3a and c show the unfiltered 2D+t and projection-based 3D+t noise series. In these global-translation benchmarks, phase correlation and rigid NoRMCorre produced numerically identical estimates because both reduce to FFT-based translational alignment against the same reference. Their error remained in the subpixel-to-low-pixel range over moderate noise, but increased sharply starting at a noise level of *∼*1.0, reaching mean errors of *∼*30.0 px in 2D+t at *σ* = 2.0 and *∼*27.8 px in projected 3D+t at *σ* = 5.0. StackReg-style intensity fitting degraded more gradually under severe noise and reached *∼*3.9 px and *∼*4.2 px at *σ* = 5.0 in the corresponding 2D+t and 3D+t benchmarks.

**Figure 3:**
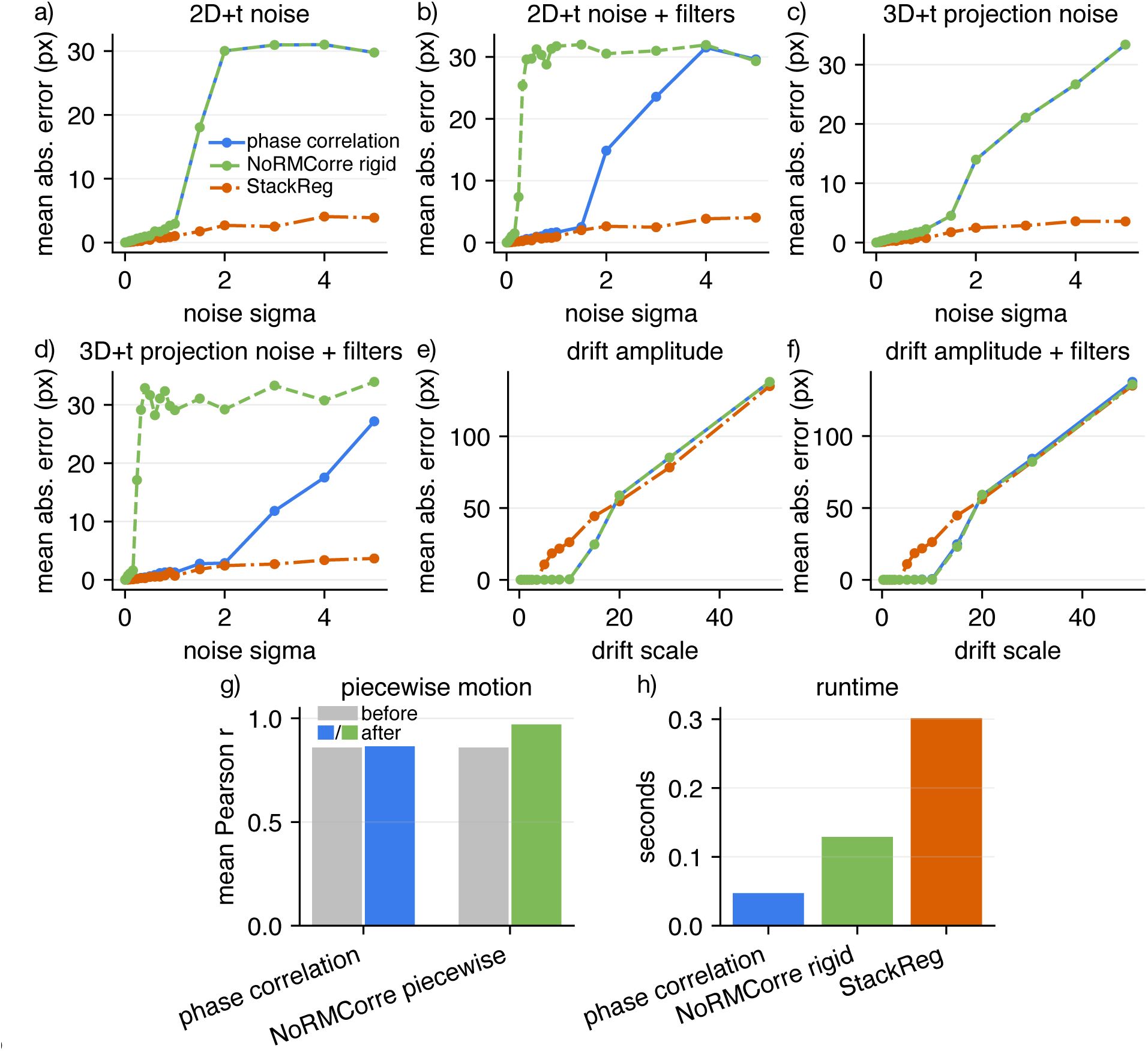
Registration robustness across noise and drift regimes. Synthetic 2D+t and 3D+t stacks contain two channels, known global translations, and progressively stronger noise or displacement amplitudes. (a,b) Translation error in 2D+t data without and with estimation prefilters. (c,d) Translation error in projection-based 3D+t data without and with estimation prefilters. (e,f) Translation error as drift amplitude increases without and with estimation prefilters. (g) Correlation before and after correction for a spatially varying local-motion benchmark. (h) Runtime comparison across representative methods. Phase correlation is plotted as a solid blue line, NoRMCorre as a dashed green line, and StackReg as a dash-dotted orange line; filtered variants keep the same method colors.

Panels b and d repeat the same experiment after estimation-only filtering. Median filtering shifted the phase-correlation failure boundary to higher noise levels: in 2D+t, phase-correlation error was 2.5 px at *σ* = 1.5 with filtering compared with 18.0 px without filtering, but it still failed at sufficiently high noise. The same trend was visible in projected 3D+t, where filtered phase correlation remained below *∼*3.6 px until *σ* = 2.0 and failed only at stronger noise. StackReg was largely insensitive to the median-filter setting in these synthetic data. In contrast, the NoRMCorre high-pass setting was detrimental for this spot-like benchmark, with errors rising already at low noise. This is consistent with a mismatch between the filtering model and the synthetic signal: the high-pass filter can attenuate broad spot structure and make the patch templates less informative, whereas it is intended primarily to suppress slowly varying background in calcium-imaging-like data.

Panels e and f probe a different failure mode: even high-SNR data become difficult when drift moves structures toward the edge of the field of view or outside the region shared with the reference. In this regime, phase correlation and rigid NoRMCorre were more tolerant of increasing displacement than StackReg. StackReg failed abruptly around a drift scale of 5, while the FFT-based estimates remained close to subpixel accuracy through drift scales of about 6.5–10 and then failed when overlap became insufficient. Estimation filtering did not rescue the large-drift regime, indicating that the dominant limitation was shared image support rather than high-frequency noise.

Panel g uses a local-motion benchmark rather than a global translation benchmark. Here the relevant quantity is not direct shift ground truth for a single global transform, but whether the corrected stack becomes more similar to the template. In this spatially varying motion field, global phase correlation produced only a small correlation increase from 0.859 to 0.865, whereas piecewise NoRMCorre increased the mean correlation to 0.971. This separates the two use cases: a single global transform is sufficient for global drift, but cannot fully describe local wave-like displacements across the field of view. Panel h reports runtime for the same implementations, because a method that is accurate but too slow for large time series may still be impractical. Together, these experiments support a practical interpretation rather than a single universal ranking: FFT-based methods are strong default choices for translational motion, StackReg can be more graceful under very noisy global-alignment conditions, and NoRMCorre-style piecewise correction becomes most relevant when the motion field truly varies spatially and each patch contains enough structure to constrain a local displacement.

### Real calcium-imaging 2D+t registration

We next applied ZenReg to a real biological calcium-imaging example movie from CaImAn [19]. The file contains 3000 frames of a single-plane functional recording and is, thus, treated as a 2D+t registration problem. Registration used all frames to construct the template, phase correlation for translational motion estimation, median filtering during estimation, and zero clipping of unsupported borders.

Figure 4 shows the same output logic as the synthetic 2D+t benchmark shown in Figure 2, now without ground-truth motion. The maximum absolute estimated correction was 2.30 px in *y* and 2.50 px in *x*, consistent with a small but visible motion artifact. The mean Pearson correlation to the template increased from 0.334 before registration to 0.487 after registration, while the registered stack was clipped laterally from 170*×*170 to 164*×*164 pixels.

**Figure 4:**
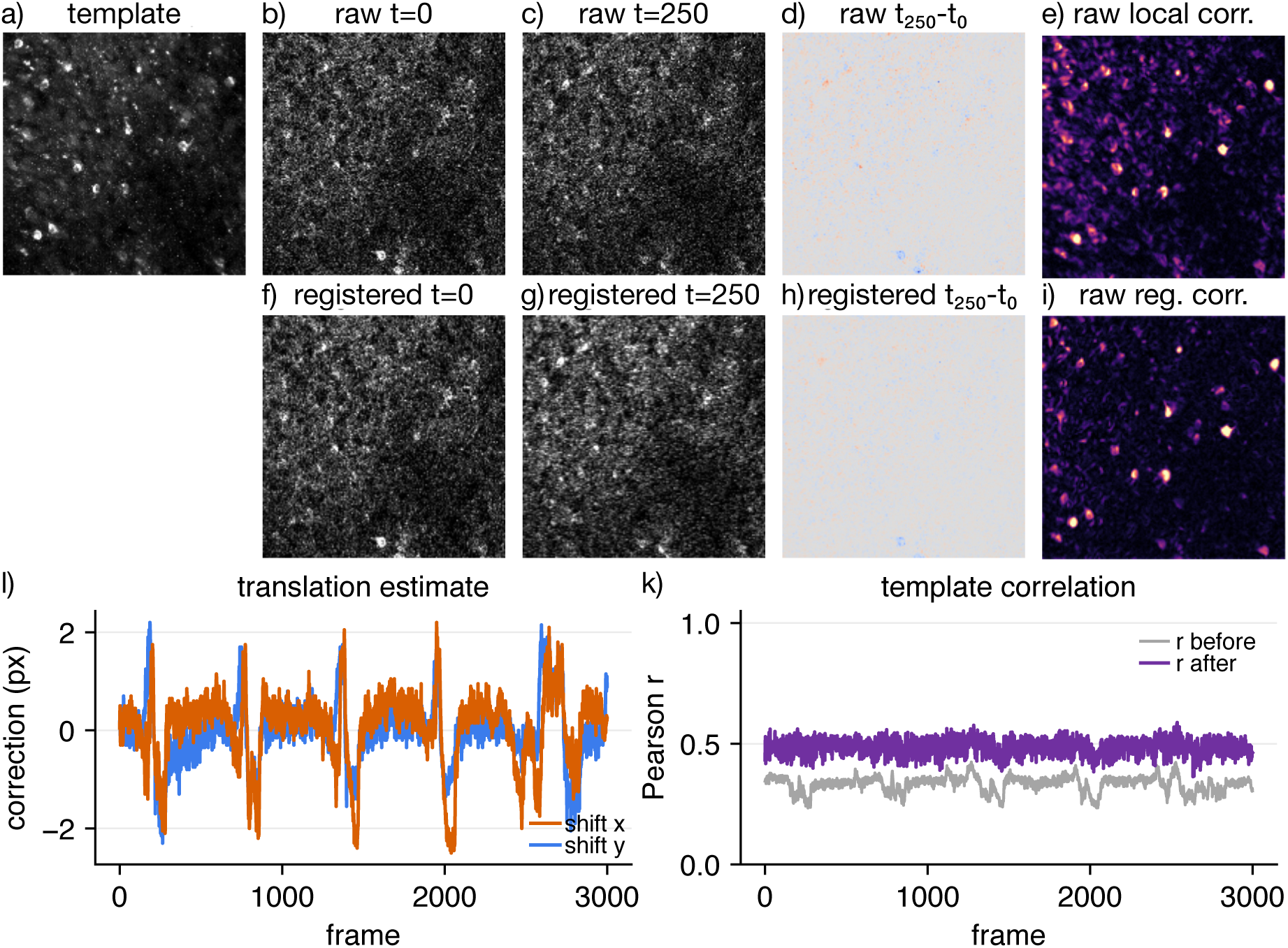
Real calcium-imaging 2D+t registration. A CaImAn example movie was registered as a 3000-frame 2D+t calcium-imaging stack. (a) All-frame registration template. (b–e) Raw data: reference frame, representative moving frame (*t* = 250), raw difference image, and local temporal correlation map. (f–i) Corresponding registered data after translational correction and zero clipping. Difference images are shown for continuity with Figure 2, but calcium activity is asynchronous across cells and time; local correlation maps are therefore more informative for this functional dataset. (j) Estimated lateral correction over time. (k) Pearson correlation to the registration template before and after correction. Data source: CaImAn demo movie [19].

For calcium imaging, raw difference images (Figure 4d and h) can be difficult to interpret because biological activity changes from frame to frame: a cell can become brighter or dimmer even when the field of view is perfectly aligned. We therefore also plotted local temporal correlation images, computed by correlating each pixel’s fluorescence time course with the time courses of its immediate spatial neighbors and averaging the resulting local correlations (Figure 4e and i). High values indicate locally coherent fluorescence dynamics, as expected for cell bodies or structured neuropil, whereas unstable background fluctuations remain weak. After registration, cell-like structures became more sharply localized in this correlation map and background-noise-like correlations were reduced (Figure 4i). This illustrates why motion correction is a practical upstream step for calcium analysis: it improves spatial consistency before downstream segmentation or source extraction, where motion-induced local correlations can otherwise contribute to false positive structures.

### Full 3D rigid registration

We then evaluated six-degree-of-freedom 3D rigid registration on two synthetic 3D+t families: dense structural volumes and sparse puncta volumes. The dense case is favorable for intensity-based Euler optimization because image information is distributed throughout the volume. The puncta case is favorable for the point backend because discrete local maxima provide a sparse geometric point cloud. We therefore treated the SimpleITK backend and the point-cloud backend as complementary rather than interchangeable methods. Each dataset was nevertheless passed to both backends.

Figure 5a–e show the dense benchmark before and after correction, using maximum projections only as a visual summary of the full volumetric transformation. Panels f–j show the corresponding puncta benchmark, where peak geometry rather than diffuse texture provides the strongest registration signal. Panels k and l quantify the actual recovered motion parameters: translation error is measured in voxels, while rotation error is measured in degrees for rotations around *z*, *y*, and *x*. In the dense structural benchmark, the SimpleITK backend recovered the imposed transform with a mean translation error of 0.018 px and a mean rotation error of 0.023 degrees across frames. The point backend was also able to operate on this feature-rich dense volume, but was less precise, reaching 0.347 px and 0.227 degrees.

**Figure 5:**
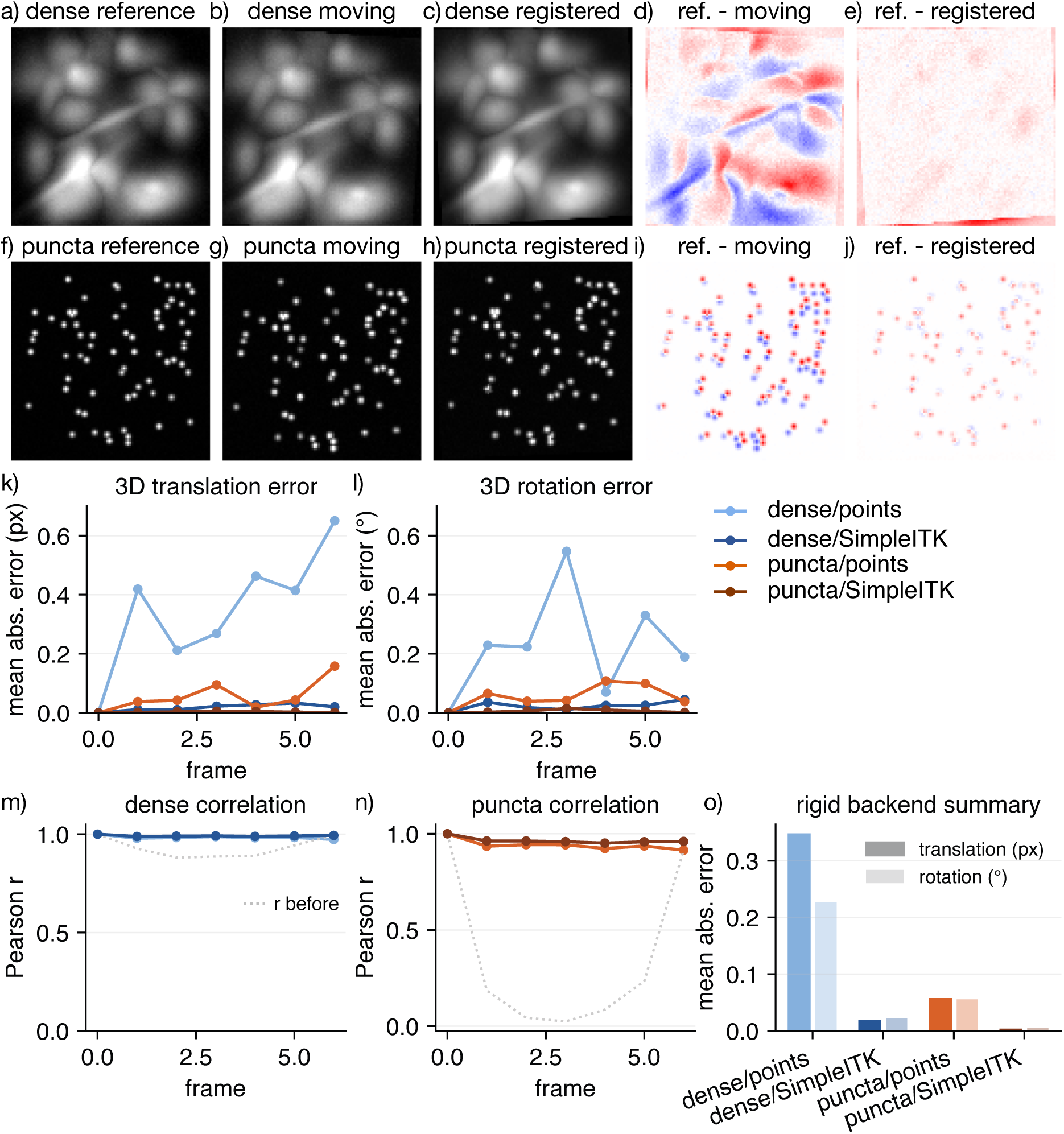
Full 3D rigid registration. Synthetic 3D+t stacks contain two channels, 20 axial planes, known translations, and known rotations around all three axes. In all cases, the transform was estimated from one channel and applied to all channels. (a–e) Dense structural benchmark: reference volume, moving volume, registered moving volume, reference–moving residual, and reference–registered residual, shown as maximum-intensity projections for visual inspection. (f–j) The same panel sequence for a sparse puncta benchmark. (k,l) Translation and rotation error across time for all backend/data combinations. (m,n) Pearson correlation to the reference before and after correction, shown separately for the dense and puncta benchmarks; grey dotted traces indicate the corresponding unregistered moving volumes. (o) Mean backend summary across frames. Dense rigid optimization uses SimpleITK/ITK [15, 14]; synthetic benchmark data were generated with ZenReg.

In the puncta benchmark, SimpleITK recovered the imposed transform with a mean translation error of 0.002 px and a mean rotation error of 0.005 degrees, while the point backend reached 0.056 px and 0.056 degrees. Panels m and n ask whether the corrected volume is more similar to the reference than the raw moving volume. For the dense benchmark, the unregistered mean Pearson correlation was 0.932 and increased to 0.992 with SimpleITK and 0.984 with the point backend. For the puncta benchmark, the unregistered mean correlation was much lower, 0.355, because sparse peaks were displaced out of overlap. After correction, it increased to 0.965 with SimpleITK and 0.942 with the point backend. Panel o summarizes the translation and rotation error families across backend/data combinations. These results support the intended interpretation of the two full-volume backends: intensity-based optimization is strongest when distributed volumetric texture is present, while point-cloud registration remains useful when enough discrete peaks can be detected and matched.

This benchmark separates two biologically distinct regimes. Dense intensity-based optimization is appropriate for structural volumes with distributed texture, while point-based alignment is appropriate for sparse puncta in which identifiable local maxima are more informative than global intensity correlations. The comparison also emphasizes the importance of physical voxel spacing. Without spacing, an anisotropic axial sampling interval would make the optimizer treat one voxel in *z* as geometrically equivalent to one pixel in *x* or *y*, leading to incorrect rotation geometry.

### Real three-photon 3D+t registration

To test the same 3D machinery on biological image content, we used a real two-channel three-photon structural volume from a published in vivo microscopy dataset [4, 30]. The source volume has no time axis, so we generated a controlled pseudo-time benchmark by extracting a central 96*×*384*×*384 voxel subvolume and applying known continuous transformations to create ten time points. A first pseudo-time series contained subpixel translations in *z, y, x*. A second series contained subpixel translations plus mild rotation around the optical axis. Transformations were estimated from channel 0 and applied to both channels.

For the translation-only pseudo-time stack, full-volume phase correlation increased mean reference correlation from 0.232 before registration to 0.720 after registration and recovered the imposed shift with a mean absolute error of 0.055 px. For the translation-plus-rotation stack, SimpleITK-based rigid registration increased mean correlation from 0.250 to 0.662, with a mean translation error of 0.068 px and a mean rotation error of 0.015 degrees. The remaining residual structure in Figure 6e and j is expected: the imposed transformations were continuous and subpixel, and the registered output is sampled back onto a discrete voxel grid. In particular, through-plane subpixel motion can leave fine residuals in maximum projections even when the estimated volume transform is accurate. We also disabled zero clipping in this benchmark to keep raw and registered stacks in the same field of view for direct metric comparison. In routine visual outputs, the user can crop unsupported borders after registration.

**Figure 6:**
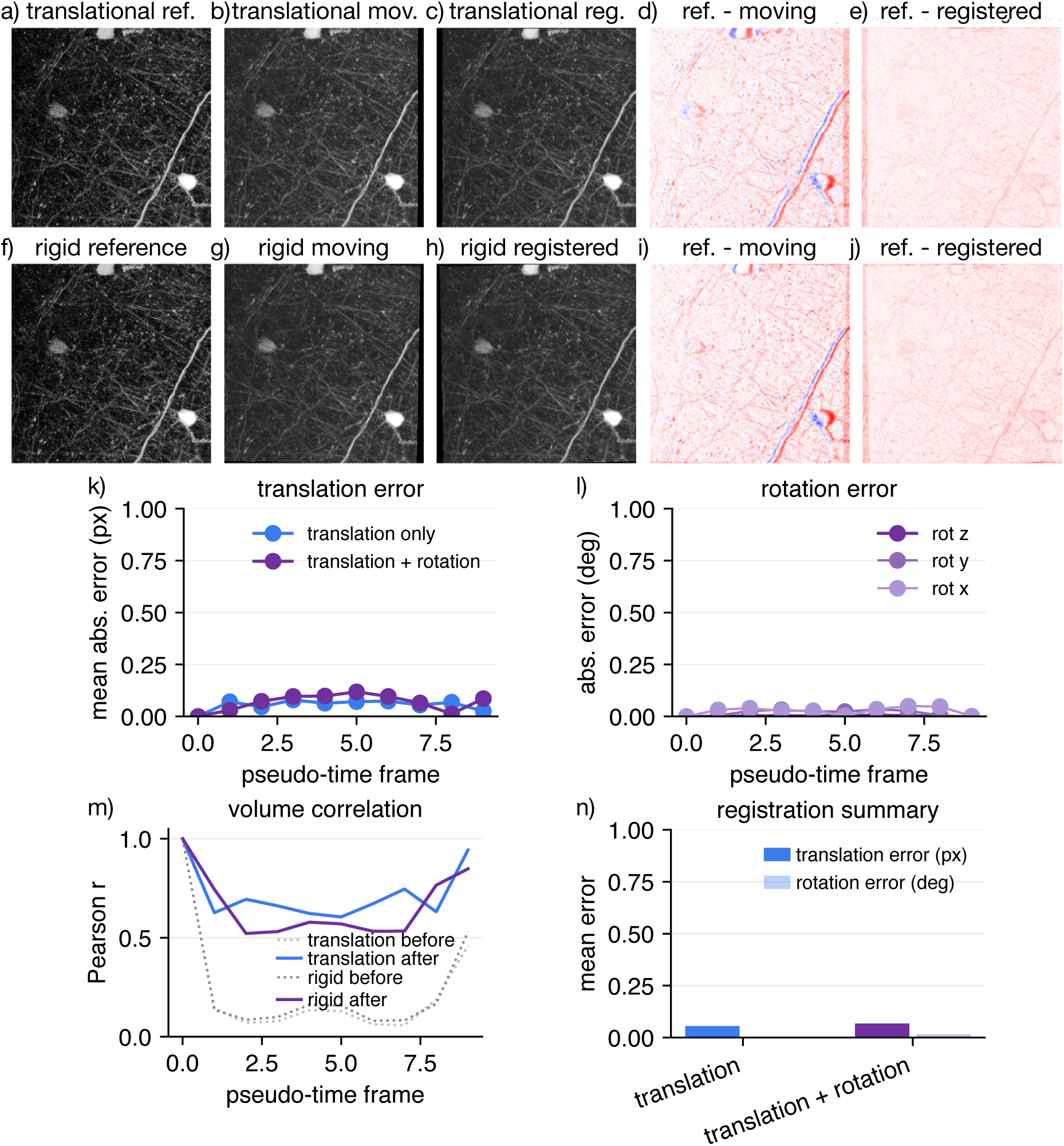
Real three-photon pseudo-time 3D registration. A real two-channel three-photon structural volume was converted into two controlled pseudo-time benchmarks. (a– e) Translation-only benchmark: reference volume, representative moving volume, registered volume, reference–moving residual, and reference–registered residual, shown as maximum-intensity projections. (f–j) Translation-plus-rotation benchmark using the same visualization sequence. (k) Translation error against the known continuous ground truth. (l) Rotation error for the SimpleITK six-degree-of-freedom rigid run. (m) Pearson correlation to the reference before and after correction. (n) ZenReg registration summary for the rigid run. Zero clipping was disabled in both examples to preserve identical shapes for metric comparison, so transform-induced zero borders remain visible in image panels. Source volume: Fuhrmann, Nebeling, and Musacchio et al. [4, 30].

### Projection-based registration and parallel execution

The choice between projection-based and full-volume registration is not only a question of accuracy, but also of computational geometry. If motion is predominantly lateral, a *z*-projection can contain enough information to estimate the relevant displacement while reducing the registration problem from a volume to an image. If axial translation is substantial, however, a projection can hide information that full 3D phase correlation can still use. We therefore added a direct benchmark comparing projection-based and full-volume translation in lateral-translation and axial-translation regimes, and a separate benchmark that repeats the same registration with different numbers of independent time points and worker processes. These benchmarks contain translations only. Rotations and six-degree-of-freedom rigid transformations are evaluated separately in Figure 5.

Figure 7a tests the regime in which projection-based registration should be sufficient: the dominant motion is lateral translation, and the projection preserves the features needed for alignment. In the full benchmark, projection-based and full-volume translation were both highly accurate in this idealized setting, with mean absolute errors of 0.021 px and 0.011 px, respectively. Panel b tests an axial-translation regime with imposed shifts in *z, y, x* but no rotation. For the chosen synthetic data, projection-based axial estimation and full-volume 3D translation again produced nearly indistinguishable errors, 0.046 px and 0.049 px, indicating that the orthogonal projections retained enough information to estimate the imposed axial shifts.

**Figure 7:**
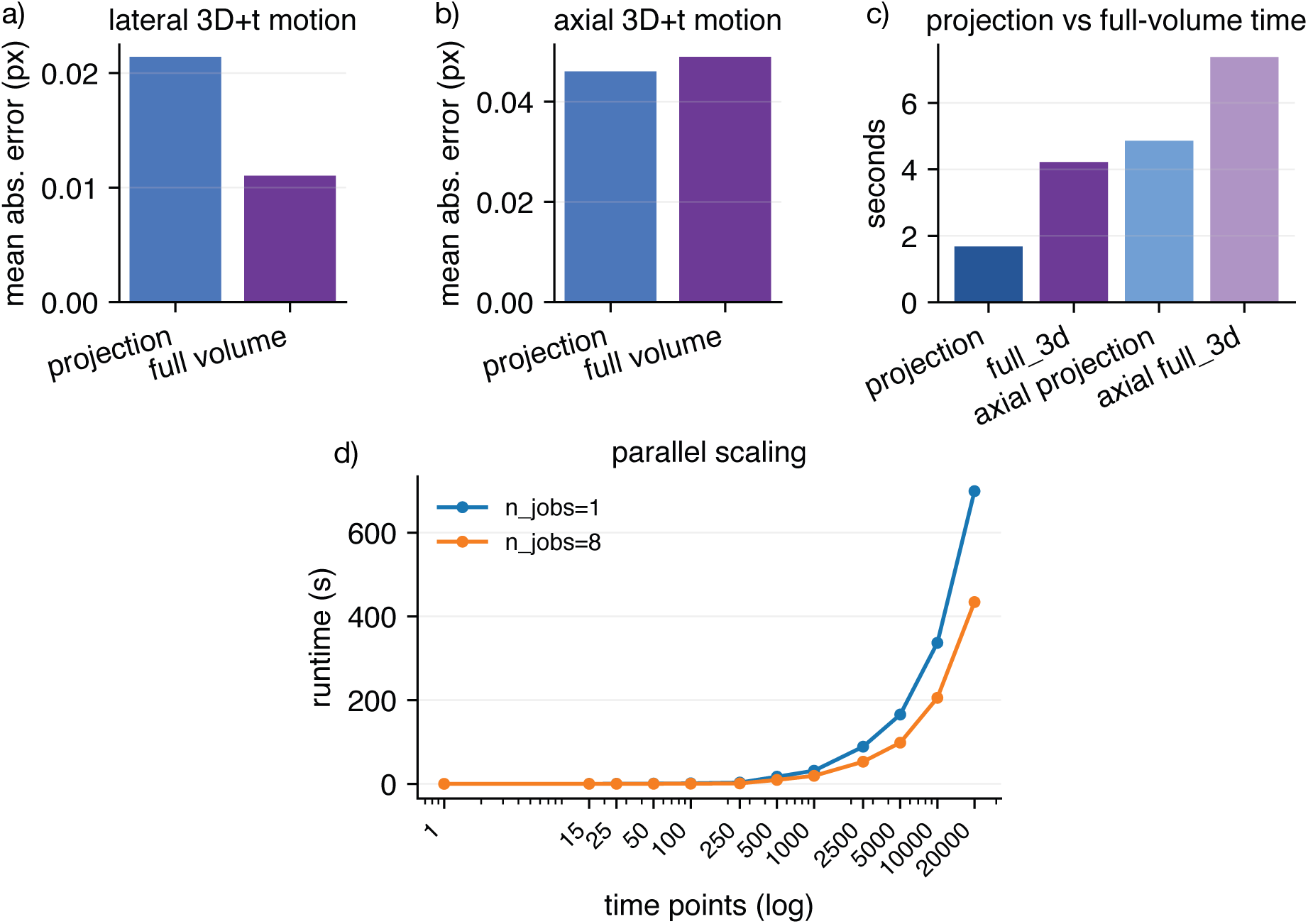
Projection-based versus full-volume registration and parallel scaling. (a) Translation error for lateral 3D+t translation with projection-based and full-volume phase correlation. (b) Translation error for axial 3D+t translation. Both panels contain translational motion only and no rotation. Here, full volume denotes full 3D translational phase correlation in *z, y, x* and not six-degree-of-freedom rigid registration. (c) Runtime comparison for the same projection/full-volume workflows. (d) Runtime as a function of time-series length using one or eight worker processes. The full benchmark samples *T* = 1, 15, 25, 50, 100, 250, 500, 1000, 2500, 5000, 10000, and 20000; quick runs use the subset up to *T* = 1000. Panel d was computed on an Apple MacBook Pro 14-inch with an M1 Max processor and 64 GB RAM.

Panel c shows why the approximation remains useful despite this similar accuracy: projection-based registration was faster in both regimes, taking 1.68 s versus 4.22 s for lateral motion and 4.86 s versus 7.38 s for axial motion. This does not imply that projections are always equivalent to full-volume registration. Rather, it identifies a favorable regime in which the faster approximation is sufficient. Stacks with complex axial structure, large z-dependent appearance changes, or rotations can break this approximation and require full-volume translation or full rigid-volume registration.

Panel d quantifies the practical benefit of parallelizing independent time points across a broad range of time-series lengths. The *T* = 1 point is a no-op baseline, included to anchor the curve at minimal time-series length. For very short sequences the absolute runtime is small regardless of worker count, so parallelization has little practical consequence. For longer time series, the savings become substantial: at *T* = 500, the runtime decreased from 17.1 s with one worker to 9.4 s with eight workers, and at *T* = 20000 from 699.0 s to 434.1 s. Together, these panels support a workflow recommendation: use projection-based registration when the biological and acquisition geometry make axial motion negligible, reserve full-volume registration for data in which axial displacement is part of the motion model, and distribute independent frames over multiple workers when time-series length makes the absolute runtime meaningful.

### Memory-efficient execution

We then evaluated ZenReg’s memory behavior on a 1.0 GB synthetic time series written as OME-TIFF and processed either through OMIO disk-backed Zarr caches or as an ordinary in-memory array. To separate registration-related allocation from the Python interpreter and imported libraries, the profiling workflow records resident set size and unique set size as changes relative to the process baseline during loading, registration, and saving. Resident set size includes all memory pages mapped into the process, including shared and file-backed pages, whereas unique set size is the private memory that would be released if the process ended. The benchmark is executed in separate processes for cold cache construction, warm cache reuse, and ordinary in-memory processing. This benchmark captures the infrastructure problem that motivates ZenReg: a registration platform should not require multiple full-size in-memory copies of a large microscopy stack.

The intended use case is common in microscopy laboratories: data are acquired to a server or network-attached storage, copied or cached once to a local scratch disk, registered while accessing only the chunks needed by the current processing step, and then written back as an OME-compatible result. Figure 8a illustrates this workflow. Panels b and c show how memory evolves over the actual load–register–save sequence for memory-mapped and in-memory execution after subtracting each process baseline. In the warm-cache memory-mapped run, peak additional RSS was 1.05 GB and peak additional USS was 0.36 GB. The corresponding in-memory run reached 2.38 GB additional RSS and 2.16 GB additional USS. Thus, the disk-backed workflow reduced private memory allocation by approximately sixfold in this benchmark, while also avoiding repeated reconstruction of the local cache when a validated cache was already present. The cold-cache run had a similar peak private-memory footprint to the warm-cache run (0.33 GB USS), but included the initial cost of constructing the disk-backed cache.

**Figure 8:**
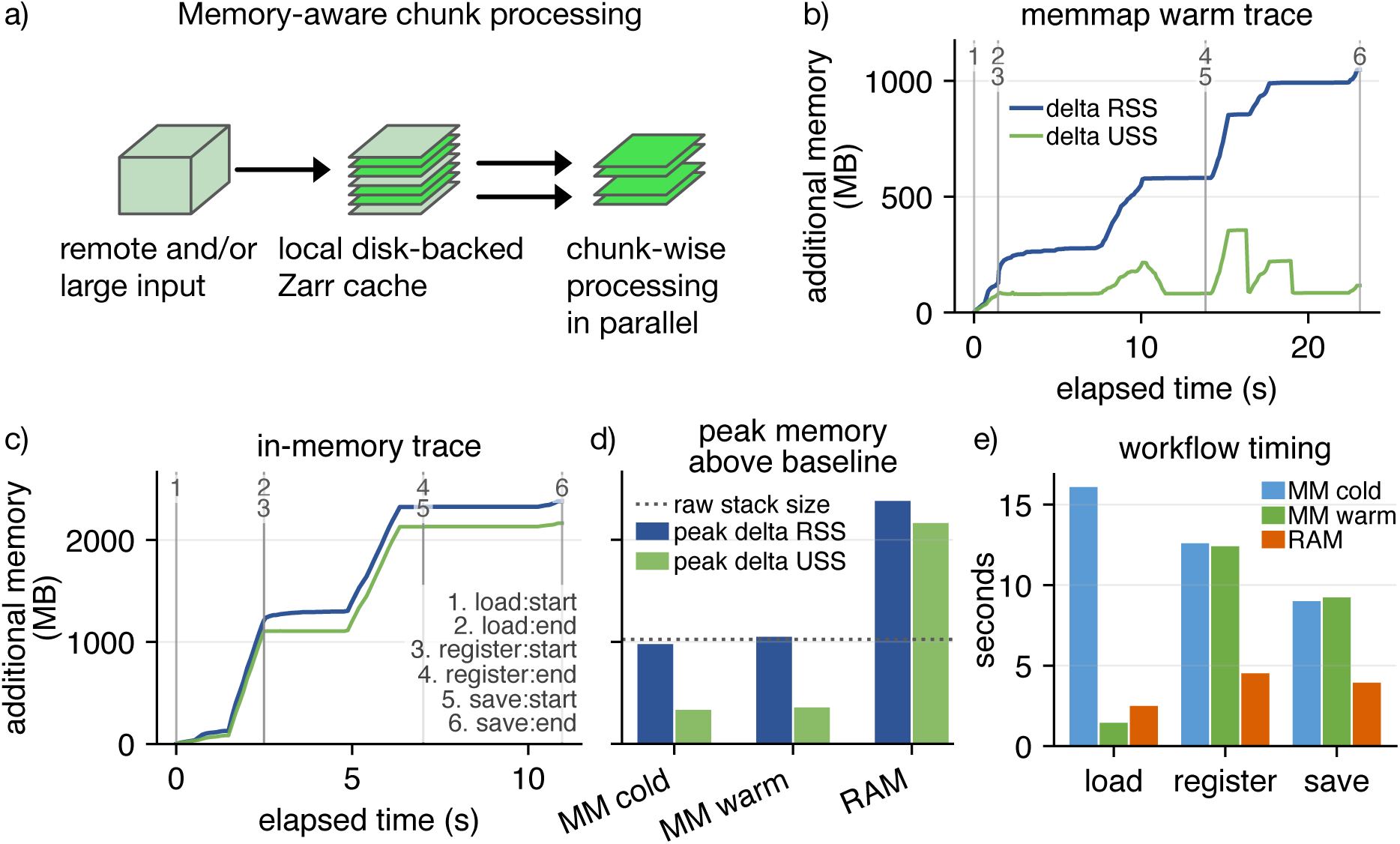
Memory-aware registration workflow. (a) Remote or large microscopy inputs can be materialized once as a local disk-backed Zarr cache, after which ZenReg accesses only the chunks required by the current step and parallelizes independent chunk-wise work where the registration model permits it. (b) Baseline-subtracted memory trace across loading, registration, and saving for disk-backed memory mapping with a reusable warm cache. (c) Corresponding baseline-subtracted trace for ordinary in-memory processing. In both trace panels, resident set size (RSS) and unique set size (USS) are shown in megabytes above the process baseline; missing or isolated invalid USS samples are left as gaps. Vertical markers indicate numbered processing events. (d) Peak additional RSS and USS for cold cache construction, warm cache reuse, and ordinary in-memory processing; the dashed line shows the raw stack size. (e) Module-level timing for loading, registration, and saving in the same three workflows.

The memory savings came with an expected runtime tradeoff. The complete warm-cache memory-mapped workflow took 23.1 s, compared with 11.0 s for the in-memory workflow, because registration and saving operated through chunked disk-backed arrays rather than a contiguous in-memory array. Cold-cache construction increased total runtime to 37.7 s, dominated by loading and cache creation. However, cache reuse changed the loading step from 16.1 s in the cold-cache run to 1.4 s in the warm-cache run. This behavior is useful in exploratory registration: once a large or remote file has been locally cached, failed runs or parameter sweeps can reuse the validated cache rather than reading and converting the source file again.

## Discussion

ZenReg frames microscopy image registration as both an estimation problem and an execution problem. The estimation problem is mathematical: given a reference, a moving image, and a restricted transformation family, find the geometry-preserving transform that best aligns the registration signal. The execution problem is practical: microscopy files are heterogeneous, multidimensional, large, and often remote. Results must preserve metadata and must be auditable after the fact. ZenReg addresses both levels within one platform.

The most conservative and broadly useful component is phase-correlation translation. It is fast, mathematically grounded in the Fourier shift theorem, and supported by established subpixel refinements [10, 11]. The same formulation applies to 2D+t time series, projected 3D+t stacks, and full 3D translations by changing the dimensionality of the images being correlated. The benchmark results support this role as a default translational estimator: phase correlation recovered global shifts with subpixel accuracy in moderate-noise and moderate-drift regimes. Its failures were abrupt rather than gradual when noise or displacement destroyed enough shared image information. StackReg-style intensity fitting behaved differently, degrading more gracefully under severe noise but failing earlier in the large-drift benchmark. This distinction is useful in practice because it turns backend choice into a model- and data-matching decision rather than a universal ranking.

Projection-based workflows provide strong computational savings when lateral motion dominates, whereas full-volume registration is available when axial motion cannot be reduced to projection geometry. In the favorable synthetic regimes tested here, projection-based and full-volume translational registration were similarly accurate, but projection-based execution was faster. This should not be read as evidence that projections are always sufficient. Instead, it identifies the regime in which the approximation is justified: axial content must remain stable enough that the projection preserves the alignment information. Complex axial rearrangements, rotations, or z-dependent structures still require full-volume translation or full rigid-volume registration.

The NoRMCorre-style implementation extends ZenReg to local translation fields and is particularly relevant for calcium imaging, where nonuniform motion can be approximated by overlapping patch shifts [13, 19]. The benchmarks emphasize that this is a model-matching advantage rather than a universal accuracy advantage. Rigid NoRMCorre converges to the global translation baseline for global drift, while piecewise NoRMCorre improved the local-motion benchmark because different regions of the field of view moved differently. This also clarifies a limitation: patch-wise translations can approximate smoothly varying local displacements, but they should not be interpreted as a general rotation or deformation engine.

Full 3D rigid registration extends ZenReg beyond planar correction by estimating translations and rotations in physical volume coordinates. The dense SimpleITK backend performed best when distributed volumetric texture was present, and the point backend remained useful when sufficiently many puncta could be detected and matched. Both backends therefore solve the same six-degree-of-freedom model, but they rely on different image evidence: continuous intensity information for dense registration and detectable local maxima for point-based registration.

The two biological examples bridge these controlled benchmarks with realistic image statistics. The calcium-imaging movie shows that even modest lateral motion can reduce spatial coherence in functional recordings, and that local correlation maps can be more interpretable than frame differences when activity is asynchronous. The three-photon benchmark demonstrates that ZenReg can reconstruct both translational and rotational motion with subpixel and subdegree accuracy in real structured volumetric microscopy data, even when the imposed motion is continuous rather than pixel-discrete.

Microscopy laboratories increasingly need registration workflows for tens of gigabytes per acquisition, often acquired or stored on remote file systems. ZenReg addresses this practical constraint by using OMIO to convert heterogeneous microscope files to a canonical axis model and, when requested, to disk-backed chunked arrays. The memory benchmark showed the expected tradeoff: disk-backed execution reduced private memory allocation substantially, especially when quantified by unique set size, but was slower than ordinary in-memory execution on the 1.0 GB benchmark stack. This is a deliberate workflow option rather than a universal default. When a stack fits comfortably in memory and speed is the only goal, in-memory execution remains attractive. When memory pressure, repeated parameter tuning, or remote storage access dominates, local cache reuse becomes more important than raw single-run speed.

Parallel execution complements the memory model but solves a different bottleneck. The scaling benchmark showed little practical benefit for short sequences because the absolute runtime was already small, whereas long time series benefited from distributing independent frames across workers. This suggests a simple operational rule: memory mapping addresses feasibility and I/O locality, while parallelization addresses throughput once there are enough independent registration units to amortize scheduling overhead. The same design supports reproducibility: every registered output can be accompanied by a motion table, similarity trace, summary plot, and settings file.

Several limitations follow directly from these model choices. ZenReg does not claim to solve arbitrary nonrigid deformation. NoRMCorre-style piecewise translations can approximate local motion fields, but they are not a substitute for explicit optical-flow, biomechanical, or fully deformable registration models. Rotation around the optical axis can be estimated from polar-transformed projections, but true 3D rotation requires full rigid-volume registration and enough volumetric structure to constrain the optimizer. Sparse point-based registration depends on detectable and matchable puncta. Dense SimpleITK-based registration depends on sufficient intensity structure and appropriate physical spacing. Finally, memory mapping reduces memory pressure after cache construction, but the initial conversion cost still depends on the input format and the capabilities of the underlying reader.

Taken together, ZenReg provides a modular, memory-efficient, and reproducible platform for common N-dimensional microscopy registration tasks. By combining canonical microscopy axes, metadata-aware I/O, projection and full-volume registration, NoRMCorre-style local translation fields, 3D rigid backends, parallel execution, and standardized report sidecars, it lowers the practical barrier between mathematical registration models and routine bioimage-analysis workflows. Its main contribution is therefore not a single new registration metric, but a coherent and extensible registration layer for microscopy data that is fast enough for large stacks, transparent enough for scientific review, and structured enough for community extension.

## Data and code availability

ZenReg is available from GitHub at https://github.com/FabrizioMusacchio/ZenReg and archived on Zenodo [31]. The preprint benchmark scripts in this repository generate the synthetic datasets, benchmark tables, and figure panels used in this manuscript draft. Synthetic data are generated programmatically with stored ground truth and are not required to be committed to the repository. The calcium-imaging and three-photon example files are not redistributed with ZenReg. The repository contains download instructions and source acknowledgments for both datasets. An extensive documentation website is available at https://zenreg.readthedocs.io.

## Acknowledgments

The authors thank Sue Koay and David Tank for the CaImAn demo movie distributed with the CaImAn project [19]. The biological three-photon example uses a stack from the dataset accompanying Fuhrmann, Nebeling, and Musacchio et al. [4, 30]. The authors also thank the open-source bioimage-analysis community whose tools and standards make reproducible microscopy workflows possible. ZenReg builds on and acknowledges OME and Bio-Formats standards [24, 25], NumPy [32], SciPy [33], scikit-image [18], Matplotlib [34], StackReg/TurboReg concepts [12], NoRMCorre and CaImAn [13, 19], SimpleITK/ITK [15, 14], OMIO [22, 23], and the broader Python scientific-computing ecosystem.

## Use of AI-assisted tools

The authors used OpenAI ChatGPT (GPT-5) to assist with language editing and wording refinement. The tools were not used to generate primary research data, perform unsupervised interpretation of results, or make authorship-level scientific decisions. All AI-assisted text changes were reviewed, verified, and edited by the authors, who take full responsibility for the accuracy, integrity, and content of the submitted article.

## Author contributions

F.M. conceived and implemented ZenReg, designed the benchmarks, and wrote the manuscript draft. M.F. supervised the project.

## References

[1] D. S. Greenberg and J. N. D. Kerr. “Automated correction of fast motion artifacts for two-photon imaging of awake animals.” In: Journal of Neuroscience Methods 176.1 (2009), pp. 1–15. doi: 10.1016/j.jneumeth.2008.08.020.

[2] R. Hattori and T. Komiyama. “PatchWarp: Corrections of non-uniform image distortions in two-photon calcium imaging data by patchwork affine transformations.” In: Cell Reports Methods 2.4 (2022), p. 100205. doi: 10.1016/j.crmeth.2022.100205.

[3] T. Wang, D. G. Ouzounov, C. Wu, N. G. Horton, B. Zhang, C.-H. Wu, Y. Zhang, M. J. Schnitzer, and C. Xu. “Three-photon imaging of mouse brain structure and function through the intact skull.” In: Nature Methods 15.10 (2018), pp. 789–792. doi: 10.1038/s41592-018-0115-y.

[4] F. Fuhrmann, F. C. Nebeling, F. Musacchio, M. Mittag, S. Poll, M. Müller, E. A. Giovannetti, M. Maibach, B. Schaffran, E. R. Burnside, I. C. W. Chan, A. S. Lagurin, N. Reichenbach, S. Kaushalya, H. Fried, S. Linden, G. C. Petzold, G. Tavosanis, F. Bradke, and M. Fuhrmann. “Three-photon in vivo imaging of neurons and glia in the medial prefrontal cortex with sub-cellular resolution.” In: Communications Biology 8 (2025), p. 795. doi: 10.1038/s42003-025-08079-8.

[5] J. B. A. Maintz and M. A. Viergever. “A survey of medical image registration.” In: Medical Image Analysis 2.1 (1998), pp. 1–36. doi: 10.1016/S1361-8415(01)80026-8.

[6] J. P. W. Pluim, J. B. A. Maintz, and M. A. Viergever. “Mutual-information-based registration of medical images: a survey.” In: IEEE Transactions on Medical Imaging 22.8 (2003), pp. 986–1004. doi: 10.1109/TMI.2003.815867.

[7] F. Helmchen and W. Denk. “Deep tissue two-photon microscopy.” In: Nature Methods 2 (2005), pp. 932–940. doi: 10.1038/nmeth818.

[8] C. Grienberger and A. Konnerth. “Imaging calcium in neurons.” In: Neuron 73.5 (2012), pp. 862–885. doi: 10.1016/j.neuron.2012.02.011.

[9] M. Pachitariu, C. Stringer, M. Dipoppa, S. Schröder, L. F. Rossi, H. Dalgleish, M. Carandini, and K. D. Harris. “Suite2p: beyond 10,000 neurons with standard two-photon microscopy.” In: bioRxiv (2017), p. 061507. doi: 10.1101/061507.

[10] C. D. Kuglin and D. C. Hines. “The phase correlation image alignment method.” In: Proceedings of the IEEE International Conference on Cybernetics and Society. 1975, pp. 163–165.

[11] M. Guizar-Sicairos, S. T. Thurman, and J. R. Fienup. “Efficient subpixel image registration algorithms.” In: Optics Letters 33.2 (2008), pp. 156–158. doi: 10.1364/OL.33.000156.

[12] P. Thévenaz, U. E. Ruttimann, and M. Unser. “A pyramid approach to subpixel registration based on intensity.” In: IEEE Transactions on Image Processing 7.1 (1998), pp. 27–41. doi: 10.1109/83.650848.

[13] E. A. Pnevmatikakis and A. Giovannucci. “NoRMCorre: An online algorithm for piecewise rigid motion correction of calcium imaging data.” In: Journal of Neuroscience Methods 291 (2017), pp. 83–94. doi: 10.1016/j.jneumeth.2017.07.031.

[14] T. S. Yoo, M. J. Ackerman, W. E. Lorensen, W. Schroeder, V. Chalana, S. Aylward, D. Metaxas, and R. Whitaker. “Engineering and algorithm design for an image processing API: a technical report on ITK–the Insight Toolkit.” In: Studies in Health Technology and Informatics 85 (2002), pp. 586–592. doi: 10.3233/978-1-60750-929-5-586.

[15] B. C. Lowekamp, D. T. Chen, L. Ibáñez, and D. Blezek. “The Design of SimpleITK.” In: Frontiers in Neuroinformatics 7 (2013), p. 45. doi: 10.3389/fninf.2013.00045.

[16] J. Schindelin, I. Arganda-Carreras, E. Frise, V. Kaynig, M. Longair, T. Pietzsch, S. Preibisch, C. Rueden, S. Saalfeld, B. Schmid, J.-Y. Tinevez, D. J. White, V. Hartenstein, K. Eliceiri, P. Tomancak, and A. Cardona. “Fiji: an open-source platform for biological-image analysis.” In: Nature Methods 9 (2012), pp. 676–682. doi: 10.1038/nmeth.2019.

[17] C. T. Rueden, J. Schindelin, M. C. Hiner, B. E. DeZonia, A. E. Walter, E. T. Arena, and K. W. Eliceiri. “ImageJ2: ImageJ for the next generation of scientific image data.” In: BMC Bioinformatics 18 (2017), p. 529. doi: 10.1186/s12859-017-1934-z.

[18] S. van der Walt, J. L. Schönberger, J. Nunez-Iglesias, F. Boulogne, J. D. Warner, N. Yager, E. Gouillart, T. Yu, and the scikit-image contributors. “scikit-image: image processing in Python.” In: PeerJ 2 (2014), e453. doi: 10.7717/peerj.453.

[19] A. Giovannucci, J. Friedrich, P. Gunn, J. Kalfon, B. L. Brown, S. A. Koay, J. Taxidis, F. Najafi, J. L. Gauthier, P. Zhou, B. S. Khakh, D. W. Tank, D. B. Chklovskii, and E. A. Pnevmatikakis. “CaImAn an open source tool for scalable calcium imaging data analysis.” In: eLife 8 (2019), e38173. doi: 10.7554/eLife.38173.

[20] S. Klein, M. Staring, K. Murphy, M. A. Viergever, and J. P. W. Pluim. “elastix: A toolbox for intensity-based medical image registration.” In: IEEE Transactions on Medical Imaging 29.1 (2010), pp. 196–205. doi: 10.1109/TMI.2009.2035616.

[21] M. D. Wilkinson et al. “The FAIR Guiding Principles for scientific data management and stewardship.” In: Scientific Data 3 (2016), p. 160018. doi: 10.1038/sdata. 2016.18.

[22] F. Musacchio, H. Antony, A. Baijal, S. Crux, F. Fuhrmann, N. Gockel, D. M. Hoffmann, D. Mercan, F. C. Nebeling, K. Wolff, and M. Fuhrmann. “OMIO: A policy-driven Python library for reproducible microscopy image I/O.” In: bioRxiv (2026). bioRxiv 2026.06.09.731118. doi: 10.64898/2026.06.09.731118.

[23] F. Musacchio. OMIO: A policy-driven Python library for reproducible microscopy image I/O. Version 0.2.6. 2025. doi: 10.5281/zenodo.18030883.

[24] I. G. Goldberg, C. Allan, J.-M. Burel, D. Creager, A. Falconi, H. Hochheiser, J. Johnston, J. Mellen, P. K. Sorger, and J. R. Swedlow. “The Open Microscopy Environment (OME) Data Model and XML file: open tools for informatics and quantitative analysis in biological imaging.” In: Genome Biology 6.5 (2005), R47. doi: 10.1186/gb-2005-6-5-r47.

[25] M. Linkert, C. T. Rueden, C. Allan, J.-M. Burel, W. Moore, A. Patterson, B. Loranger, J. Moore, C. Neves, D. MacDonald, A. Tarkowska, C. Sticco, E. Hill, M. Rossner, K. W. Eliceiri, and J. R. Swedlow. “Metadata matters: access to image data in the real world.” In: Journal of Cell Biology 189.5 (2010), pp. 777–782. doi: 10.1083/jcb.201004104.

[26] W. Kabsch. “A solution for the best rotation to relate two sets of vectors.” In: Acta Crystallographica Section A 32.5 (1976), pp. 922–923. doi: 10.1107/S0567739476001873.

[27] M. A. Fischler and R. C. Bolles. “Random sample consensus: a paradigm for model fitting with applications to image analysis and automated cartography.” In: Communications of the ACM 24.6 (1981), pp. 381–395. doi: 10.1145/358669.358692.

[28] R. C. Gonzalez and R. E. Woods. Digital Image Processing. 4th ed. Pearson, 2018. isbn: 978-1-292-22304-9.

[29] H. Hwang and R. A. Haddad. “Adaptive median filters: new algorithms and results.” In: IEEE Transactions on Image Processing 4.4 (1995), pp. 499–502. doi: 10.1109/83.370679.

[30] F. Fuhrmann, F. C. Nebeling, F. Musacchio, et al. Data from: Three-photon in vivo imaging of neurons and glia in the medial prefrontal cortex with sub-cellular resolution. Dataset. 2025. doi: 10.5061/dryad.tqjq2bw90.

[31] F. Musacchio. ZenReg: Fast and memory-efficient N-dimensional microscopy image registration for Python. 2026. doi: 10.5281/zenodo.21727826.

[32] C. R. Harris et al. “Array programming with NumPy.” In: Nature 585 (2020), pp. 357–362. doi: 10.1038/s41586-020-2649-2.

[33] P. Virtanen et al. “SciPy 1.0: fundamental algorithms for scientific computing in Python.” In: Nature Methods 17 (2020), pp. 261–272. doi: 10.1038/s41592-019-0686-2.

[34] J. D. Hunter. “Matplotlib: A 2D Graphics Environment.” In: Computing in Science & Engineering 9.3 (2007), pp. 90–95. doi: 10.1109/MCSE.2007.55.

